# SEME enables programmable transcript reconstruction through synthetic microexon incorporation

**DOI:** 10.64898/2026.09.13.749216

**Authors:** Myeongjune Go, Jaewon Oh, Seunghee Moon, Elaine Zhelan Chen, Byunggik Kim, David Suh, Sangwoo Kim, Chulan Kwon, Seunghyun Lee

**Affiliations:** Division of Cardiology, Department of Medicine, Johns Hopkins University School of Medicine; Baltimore, MD 21205, USA; Division of Cardiology, Severance Cardiovascular Hospital, Cardiovascular Research Institute, Yonsei University College of Medicine; Seoul 03722, Republic of Korea; Department of Biomedical Systems Informatics and Brain Korea 21 PLUS Project for Medical Science, Yonsei University College of Medicine; Seoul 03722, Republic of Korea

## Abstract

Precise correction of pathogenic mutations remains challenging for therapeutic applications because genome editing within protein-coding exons can generate unintended insertion–deletion byproducts that disrupt coding integrity and protein function. To address this limitation, we developed the Spliceable Editable Microexon Element (SEME), an intron-targeting platform that enables programmable microexon incorporation through endogenous splicing. In induced pluripotent stem cell-derived cardiomyocytes (iPSC-CMs), SEME corrected aberrant splicing caused by the dilated cardiomyopathy-associated FLNC c.2003A>G variant by restoring the five nucleotides missing from exon 12, thereby recovering filamin C expression. Together, these findings establish programmable microexon incorporation as a proof-of-concept strategy for correcting diverse disease-causing transcript defects.

## Introduction

Pathogenic variants that disrupt transcript integrity are major causes of human disease, including cardiomyopathies, neuromuscular disorders, cancer, and neurological diseases (*1-4*). Many of these variants generate out-of-frame transcripts that result in truncated protein synthesis or nonsense-mediated mRNA decay (NMD). Advances in next-generation sequencing have accelerated the identification of disease-associated splice-altering and frameshift variants, revealing that defects affecting RNA processing and translational reading frames are widespread contributors to human pathology (*5, 6*). Despite substantial progress in genome-engineering technologies, therapeutic correction of these mutations remains challenging. Many gene-editing strategies directly modify protein-coding exons to correct pathogenic sequences or translational reading frames. However, repair of nuclease-induced DNA double-strand breaks can generate heterogeneous on-target outcomes, including small insertions and deletions (indels) as well as larger indels and complex genomic rearrangements. When these unintended byproducts occur within coding exons, they may disrupt the translational reading frame or compromise protein function (*7, 8*). However, genome editing can also generate unintended on-target alterations, including small insertions and deletions, larger deletions, and, particularly with nuclease-based approaches, complex genomic rearrangements (*9, 10*). When these unintended byproducts occur within coding exons, they may disrupt the translational reading frame or compromise protein function.

To circumvent these limitations, therapeutic approaches such as exon skipping have been developed to restore transcript continuity at either the genomic level through CRISPR–Cas9-mediated exon disruption or the RNA level through antisense oligonucleotide (ASO)-mediated splicing modulation. These strategies are being extensively investigated for Duchenne muscular dystrophy (DMD) and related genetic disorders in which restoration of the translational reading frame can be achieved despite partial exon loss (*11-14*). However, exon skipping necessarily removes one or more coding exons, and, for some mutations, restoration of the reading frame requires exclusion of additional exons beyond the mutation-containing exon. The resulting loss of coding sequence may further truncate the transcript and compromise protein function when the excluded regions encode essential structural or functional domains (*15-17*). These limitations highlight the need for alternative strategies capable of reconstructing functional transcripts while minimizing direct perturbation and unnecessary loss of protein-coding sequence.

To address this challenge, we explored whether endogenous intronic splicing environments could be harnessed as programmable platforms for transcript reconstruction. We reasoned that the splice-regulatory architecture surrounding naturally recognized short exons could be repurposed to support efficient incorporation of programmable microexons. Based on this concept, we engineered a modular splicing cassette consisting of a 3′ splice-site module, a programmable microexon, and a 5′ splice-site module, which we termed the Spliceable Editable Microexon Element (SEME). Using SEME, we sought to correct heterozygous FLNC splicing defects and restore disrupted translational reading frames.

## Materials and Methods

### Study approval and human participants

The human participant component of this study, including clinical evaluation, review of medical records, and identification of the FLNC variant, was conducted at Severance Hospital in accordance with the Declaration of Helsinki. The study protocol was reviewed and approved by the Institutional Review Board of Severance Hospital (IRB No. 4-2020-0112), and written informed consent was obtained from all participants before enrollment. All participant information was de-identified before analysis.

Subsequent mechanistic experiments were performed at Johns Hopkins University using gene-edited induced pluripotent stem cell lines generated from an established control iPSC line. No identifiable participant information or patient-derived biospecimens were transferred to or analyzed at Johns Hopkins University.

### *In silico* splicing pattern prediction

To evaluate the impact of FLNC variant identification in the DCM patient on splicing, SpliceAI (version 1.3.1) was employed to predict scores for acceptor gain, acceptor loss, donor gain, and donor loss. The maximum delta score among the four categories was reported as the overall SpliceAI score. To predict alterations in splicing patterns within SEME-inserted sequences, the AlphaGenome API was utilized. A 100KB genomic segment encompassing the *FLNC* coding region was retrieved from the Ensembl server using the human reference genome. From this wild-type sequence, both variant-containing and SEME-integrated sequences were constructed. Splice sites were predicted by inputting these prepared sequences into the AlphaGenome API alongside the cardiac muscle cell ontology term. The resulting coordinates for splice donors and acceptors, along with their respective usage scores, were integrated and visualized using hg38/GENCODE v46 annotation data.

### iPSC culture and cardiomyocyte differentiation

For maintenance, iPSCs were cultured in the Essential 8 Medium (Gibco) on vitronectin (Gibco) coated plates. Cells were passaged at a 1:10 ratio every four days using ReLeSR (STEMCELL Technologies), with 10 μM Y-27632 (Tocris) added to the medium. The media was replaced with fresh E8 within 24 hours of post-passage. Cardiomyocyte differentiation was performed following previously described paper (*18*). Briefly, iPSCs were dissociated using GCDR (STEMCELL Technologies), and 8×10^5^ cells were replated into a single well of a Matrigel (Corning) coated 6-well plate. Upon reaching approximately 90% confluence, the medium was replaced with 3 ml of RPMI 1640 medium supplemented with B-27 supplement (Gibco) containing 6 μM CHIR99021 (Tocris). On day 2 and 3 of differentiation, 2 ml and 1 ml of the same medium were added, respectively. On day 4, the medium was replaced with 3 ml of RPMI 1640 supplemented with B-27 minus insulin containing 2μM C59 (Selleckchem). On day 6, the medium was replaced with 3 ml of RPMI 1640 medium supplemented with B-27 minus insulin. On day 8, upon the observation of partial beating, the medium was changed to 3 ml of RPMI 1640 medium supplemented with B-27 supplement (Gibco). On day 10, metabolic purification of the iPSC-CMs was performed through glucose deprivation using RPMI 1640 no glucose medium (Gibco) supplemented with B-27 supplement. The replating of iPSC-CMs was carried out using TrypLE Select Enzyme (10X) (Gibco) according to the manufacturer’s instructions and seeded onto Matrigel coated T75 flasks in RPMI 1640 medium supplemented with B-27, 10% KO serum (Gibco) and 10 μM Y-27632.

### Immunocytochemistry

For immunocytochemistry, cells were fixed with 4% paraformaldehyde for 20 minutes at room temperature. Subsequently, the cells were permeabilized in 1X PBS containing 0.3% Triton X-100 and blocked with 6% BSA containing permeabilization buffer for 30 minutes. Primary antibodies (Table 3) were diluted in 1X PBS containing 0.3% Triton X-100 and 0.6% BSA and incubated with the cells overnight at 4°C. Secondary antibodies were then applied in the same incubation buffer for 3 hours at room temperature. Nuclei were counterstained with Hoechst 33342 (Invitrogen) at a 10000X dilution during the final wash step. LSM710 confocal microscope (Zeiss) was used for imaging. Sarcomere length was measured using ZEN Blue software (Zeiss), and cell roundness was analyzed to quantify cell morphology using Fiji software (v1.54f; https://imagej.net/fiji). The used antibodies are listed in Table 3

### Nucleic acids preparation and Sanger sequencing

Genomic DNA was extracted from iPSCs using G-spin Total DNA Extraction Mini Kit (iNtRON Biotechnology) in accordance with the manufacturer’s instructions. Total RNA was isolated from HeLa, iPSC, and iPSC-CMs using Ribospin II (GeneAll Biotechnology). For reverse transcription, 3000 ng of total RNA was converted into cDNA libraries using PrimeScript Reverse Transcriptase (TaKaRa). Prepared DNAs were amplified by Q5 High Fidelity DNA Polymerase (NEB). Amplified DNAs were purified using the MEGAquick-spin Plus Total Fragment DNA Purification Kit (iNtRON Biotechnology) following the manufacturer’s instructions. Sanger sequencing was outsourced to Bionics Co. (Republic of Korea). All primers utilized in this study are listed in Table S1.

### Quantitative real-time PCR

Total RNA isolation and cDNA synthesis were performed as previously described. Quantitative real-time PCR was conducted using the QuantStudio 3 Real-Time PCR System (Applied Biosystems) utilizing FastStart Universal SYBR Green Master Mix (ROX) (Roche) in accordance with the manufacturer’s instructions. The specific primer sequences utilized in this study are detailed in Table S2. Relative mRNA expression levels were quantified using the 2^-ΔΔCt^ method, with *GAPDH* serving as the internal housekeeping gene for normalization.

### Immunoblot analysis

Total protein was extracted by lysing cells in an ice-cold lysis buffer consisting of 0.2% (v/v) Triton X-100, 0.3% (v/v) NP-40, 150 mM NaCl, 50 mM Tris-HCl (pH 7.4), 1 mM EDTA, 1mM EGTA, 1 mM Na_3_VO_4_, 1 mM NaF, 1 mM PMSF, and 1X Xpert Protease inhibitor Cocktail Solution (GenDEPOT). Cells were homogenized via gentle pipetting followed by pulse sonication. The resulting lysates were clarified by centrifugation at 13,000 g for 10 minutes at 4 °C. Protein concentrations in the supernatants were determined using the Pierce 660 nm Protein Assay Reagent (Thermo Scientific) according to the manufacturer’s instructions.

For immunoblotting, 15 μg of protein were separated via house made SDS-PAGE gels and immobilized onto NC membranes (Cytiva). NC membranes were blocked for 1 hour at room temperature in 1X PBS containing 1% Tween – 20 and 5 % skim milk (w/v). The membranes were then probed with specific primary antibodies 1:1000 diluted in PBST with 3% BSA overnight at 4°C. Then the secondary antibodies 1:5000 diluted in PBST with 5% skim milk attached for 1 hour at room temperature. Detailed information regarding the antibodies used is provided in Table S3. The signal was detected using Pierce ECL Western Blotting Substrate (Thermo Scientific) and visualized with the Fusion SOLO 7S imaging system (Vilber Lourmat).

### Generation of minigene splicing reporter constructs

Genomic DNA from HeLa was extracted as previously described and utilized as the template for minigene insert fragments. The SV40-EGFP backbone was constructed by the ligation of an SV40 promoter, the 5’ UTR of ALDOA, a Kozak sequence including an ATG start codon, EGFP, and an SV40 poly A signal sequence. To generate the FLNC ^WT^ minigene, a genomic region spanning from the middle of exon 11 to the middle of exon 15 (hg38: Chr7: 128,841,247 – 128,842,620) was amplified. Primers were designed with homologous overlap sequences of at least 20 nucleotides to facilitate NEBuilder HiFi DNA Assembly (NEB). The minigene backbone plasmid was linearized using SwaI (NEB) following manufacturer’s instructions. The backbone and insert were assembled by HiFi assembly following manufacturer’s instructions and were transformed into HIT Competent Cells-DH5α (Real Biotech). The FLNC ^mut^ minigene was generated via site-directed mutagenesis of the WT construct and confirmed by Sanger sequencing. To construct the spliceable microexon inserted minigene plasmids (FLNC ^SEME^ 97 bp, 215 bp, and 447bp), the FLNC ^mut^ minigene was digested with SmaI (NEB). Inserts for the 215 bp and 447 bp constructs were amplified from genomic regions containing the selected splice-regulatory sequences. The FLNC ^SEME^ constructs were finalized by incorporating the 5 bp microexon sequence via site-directed mutagenesis. The insert for the 97 bp construct was prepared by PCR amplification of synthesized oligonucleotides with overlapping sequences. Each fragment was integrated into the linearized FLNC ^mut^ plasmid via HiFi assembly. All plasmids used for transfection were purified using the NucleoBond Xtra Midi Kit (Macherey-Nagel).

### Minigene transfection and in vitro splicing assay

HeLa cells were seeded into 6-well culture plates at a density of 2×10^5^ cells per well. After 24 hours, the cells were transfected with 1 μg of the minigene plasmids using the TransIT-X2 Dynamic Delivery System (TaKaRa) according to the manufacturer’s protocol. The transfected cells were incubated for 48 hours at 37 °C in humidified atmosphere containing 5% CO2. EGFP expression was visualized using an Olympus IX 71 fluorescence microscope (Olympus) prior to cell harvesting. Total RNA extraction and cDNA synthesis were performed as previously described. To selectively amplify minigene-derived transcript and avoid interference from endogenous *FLNC* expression, PCR was conducted using a primer pair targeting the 5’ UTR of *ALDOA* and the *EGFP* sequence (Table 1). The resulting PCR products were analyzed via Sanger sequencing, and sequences were aligned using SnapGene software.

### iPSC genome editing

The wild-type iPSC line, CMC-iPSC-011, was obtained from the Korea National Institute of Health (Republic of Korea) and utilized to generate all gene-edited derivative lines. gRNA targeting specific intronic sites within *FLNC* (NM_001458.5) was designed using the Invitrogen TrueDesign Genome Editor (Invitrogen), with the highest-scoring candidates selected for synthesis. The gRNA sequences were subsequently cloned to the PX458 vector via Golden Gate assembly. To construct the double-stranded DNA donor templates for gene editing, the pTubb3-MC plasmid was linearized by digestion with SmaI and ApaI. Homology arms containing target sequences for *FLNC* (hg38: Chr7: 128,840,215 – 128,841,841) was PCR-amplified and integrated into the backbone via HiFi assembly. The final double-stranded DNA donors used for transfection were prepared via PCR amplification; the associated primers are detailed in Table 1.

For genome editing, 2×10^5^ iPSCs were co-transfected with 1 μg of the gRNA-containing PX458 vector and 1.5 μg of the donor using the Neon Transfection System (Invitrogen). Following electroporation, cells were maintained in mTeSR1 media (STEMCELL Technologies) supplemented with 10 μM Y-27632 for 24 hours, followed by a medium change to fresh mTeSR1. After an additional 24-hour incubation, cells were dissociated into a single-cell suspension and GFP-positive cells were isolated into 96-well plates using an SH800S cell sorter (Sony Biotechnology). The resulting single-cell-derived colonies were validated for targeting genomic integration via PCR amplification of the target loci followed by Sanger sequencing.

### Bulk RNA sequencing

Total RNA was isolated from day-40 wild-type, FLNC ^mut^, and FLNC ^SEME^-cardiomyocytes (three biological replicates/group) using the easy-BLUE™ Total RNA Extraction Kit (iNtRON Biotechnology) and submitted to Integrated Genomics Center (IGC) at Johns Hopkins University School of Medicine (Baltimore, MD, USA) for mRNA sequencing. FASTQ files were processed using nf-core/rnaseq v3.14.0 with GRCh38 and GENCODE v44. Transcript abundance was quantified using Salmon v1.10.1. Transcripts with ≥10 estimated counts in ≥3 samples were analyzed using DESeq2 with differentiation batch and group as covariates. Differential transcript usage was assessed using a batch-blocked transcript-versus-rest quasibinomial model, with significance defined as Benjamini–Hochberg-adjusted P < 0.05 and an absolute isoform-fraction change ≥0.10. Transcript models were assembled using StringTie v2.2.1, merged, compared with GENCODE v44 using gffcompare v0.10.4, and requantified against a common catalog. Condition-specific novel junction candidates required class j assignment, mean TPM ≥1 and TPM ≥1 in ≥2 case replicates, and TPM <0.1 in every control replicate. FLNC splicing was visualized using IGV Sashimi plots. Canonical exon 12–13 junction efficiency was calculated from uniquely mapped junction-spanning reads as 2 × J12–13/(J11–12 + J13–14). The assembled MSTRG.27911.2 sequence was extracted using gffread and aligned against FLNC-201 (ENST00000325888.13) to resolve CAG-associated boundary ambiguity and assess the inferred five-nucleotide deletion. Junction efficiencies were summarized as mean ± SD and compared using two-sided Welch t tests; transcriptome-wide P values were adjusted using the Benjamini–Hochberg method.

### Calcium transient assay

Five days prior to the assay, iPSC-CMs were seeded at a density of 2×10^5^ cells onto Matrigel-coated glass bottom dishes. The cells were maintained in RPMI 1640 medium supplemented with B-27, with the medium refreshed every 48 hours. On the day of the assay, the cells were loaded with 5 μM Fluo-4 AM (Invitrogen) and incubated for 10 minutes at 37 °C. Following incubation, the loading medium was replaced with Tyrode’s salts and cells were equilibrated for an additional 5 minutes at 37 °C. Intracellular calcium dynamics were recorded in line-scan mode using LSM 710 confocal microscope (Zeiss). Fluo-4 fluorescence was excited at 488 nm, and emission was collected at approximately 516 nm, respectively. A total of 10,000 lines were acquired over a 60-second period. Raw line-scan data were processed using ZEN Black edition (Zeiss), and fluorescence intensities were normalized using ZEN Blue edition (Zeiss). Subsequent kinetic parameters analysis was performed by excel as previous published paper (*19*).

### Statistical analysis

For the normality test, the distribution of the data was confirmed applying Shapiro-Wilk normality test. Based on the results of the normality test, for cases where the p-value≥0.05, the comparison between two groups was analyzed using Student’s t-test, and the comparison among three or more groups was analyzed using One-way ANOVA with Tukey’s multiple comparison test. Values were expressed as the mean ± SD. Statistical analysis was conducted using Prism 8 (GraphPad Software). Statistical significance was set at * *p* < 0.05, ** *p* < 0.01, *** *p* < 0.001, and **** *p* < 0.0001.

## Results

### The *FLNC* c.2003A>G variant induces aberrant splicing and haploinsufficiency via NMD

We engineered a modular cassette comprising a 3′ splice-site module, a programmable microexon sequence, and a 5′ splice-site module, which we termed SEME (Fig. 1A).

**Figure 1.**
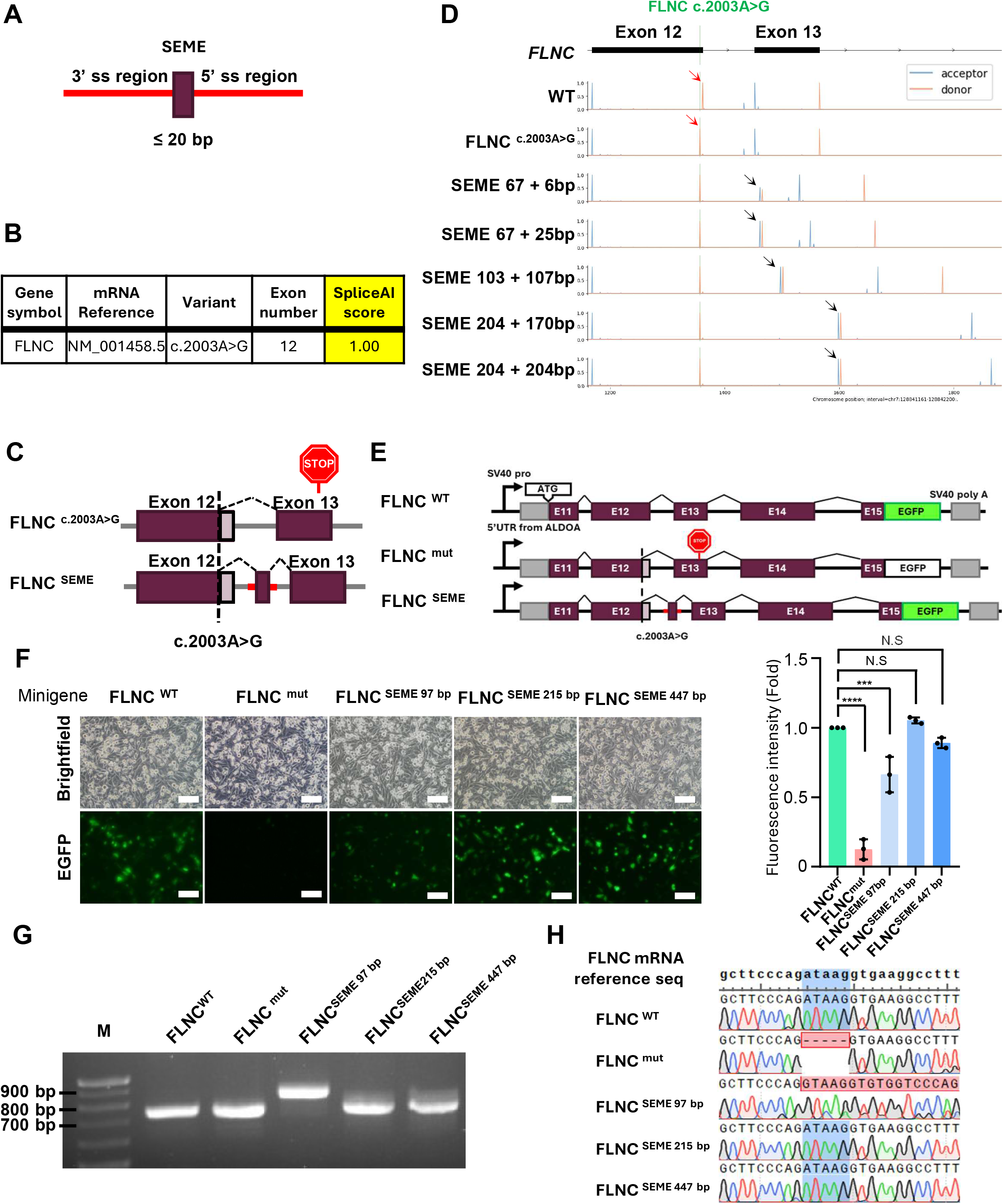
Genomic architecture of SEME and validation of *FLNC* splicing correction *in silico* and *in vitro*. (A) structural organization of the SEME. The synthetic microexon component (≤20 bp) represents the fragment integrated into the mature mRNA transcript via splice-in mechanism, while the flanking intronic sequences (red) are derived from the endogenous selected splice-regulatory sequences. (B) Identification of patient-specific *FLNC* mutations. (C) Schematic diagram of the SEME-mediated splicing correction strategy. Integration of the SEME into *FLNC* intron 12 introduces the 5 nucleotides lost due to the c.2003A>G mutation as a synthetic microexon, thereby restoring an in-frame, fully wild-type *FLNC* mRNA transcript. (D) AlphaGenome-mediated optimization of the SEME flanking sequences. Red arrows indicate the *FLNC* exon 12 splice donor site, whereas black arrows indicate the predicted splice sites of the SEME-derived microexon. (E) Schematic representation of the *FLNC* minigene reporter construct. The FLNC ^c.2003A>G^ splicing variant induces a PTC, shifting the downstream EGFP sequence out-of-frame and disrupting its translation. Conversely, FLNC ^SEME^ reconstitutes the correct reading frame, thereby successfully rescuing EGFP expression. (F) Fluorescence microscopy evaluation of minigene reporter rescue. Representative bright-field and fluorescence images of HeLa captured 48 hours post-transfection with the respective minigene constructs. Correction into an in-frame FLNC sequence drives EGFP expression, which was quantified via imageJ. Scale bar = 100μm. (G) RT-PCR analysis of minigene-derived transcripts. The expected amplicon size for WT *FLNC* is 755 bp. (H) Transcriptional validation via Sanger sequencing. Sanger sequencing chromatograms of the RT-PCR products obtained from (G). The blue-shaded region highlights the 5-nucleotide sequence typically lost due to the cryptic splicing mutation, while the red-shaded sequence denotes nucleotides that deviate from the canonical reference sequence. Data are presented as the mean ±S.D. from three independent experiments. *** *p* < 0.001, **** *p* < 0.0001 by one-way ANOVA followed by Tukey’s post-hoc test.

Among several variants of uncertain significance (VUS) identified by cardiomyopathy panel sequencing in patients with dilated cardiomyopathy from the Division of Cardiology at Severance Hospital, we identified the FLNC c.2003A>G (p.Asp668Gly) variant. SpliceAI analysis predicted that this variant strongly activates a cryptic splice site, with a maximum delta score of 1.00, suggesting a high probability of aberrant splicing (Fig. 1B). Notably, the predicted aberrant splicing event was expected to generate a premature termination codon (PTC) within exon 13. Because premature termination codons located more than 50–55 bp upstream of the final exon– exon junction commonly trigger nonsense-mediated mRNA decay (NMD), the mutant transcript was predicted to undergo NMD-mediated degradation (*20*).

### SEME-mediated microexon insertion restores the *FLNC* c.2003A>G splicing defect *in vitro*

The FLNC c.2003A>G mutation generates a cryptic splicing donor, leading to the excision of the terminal 5 bp of exon 12 and the subsequent production of a frame-shifted mRNA transcript. To counteract this, we designed a gene correction strategy involving the insertion of SEME into FLNC intron 12 to facilitate incorporation of the missing 5 bp sequence, thereby restoring the complete wild-type FLNC sequence (Fig. 1C). In this model, the SEME-mediated microexon is incorporated into the FLNC mRNA to compensate for the nucleotides lost to cryptic splicing.

In silico splicing prediction using AlphaGenome accurately predicted cryptic splice donor activation by the FLNC c.2003A>G mutation (Fig. S1), consistent with experimental observations (Fig. 1D). Extension of the SEME-associated intronic sequences progressively improved predicted splice-in efficiency.

To experimentally validate SEME-mediated splicing, we generated a minigene encompassing FLNC exons 11–15 fused to a C-terminal EGFP reporter, such that frameshifted transcripts abolished EGFP expression (Fig. 1E). The FLNC c.2003A>G mutation markedly reduced EGFP fluorescence, whereas SEME insertion restored reporter expression. However, contrary to in silico predictions, the 97 bp SEME construct (67-bp 3′ss + 25-bp 5′ss) failed to fully recover EGFP expression. Complete restoration was observed only with the extended 215-bp construct (103-bp 3′ss + 107-bp 5′ss) (Fig. 1F). Additionally, agarose gel analysis demonstrated that the 97-bp SEME construct generated a larger FLNC cDNA product than the other constructs, suggesting aberrant splicing associated with incomplete intron removal (Fig. 1G).

Sanger sequencing of minigene-derived cDNA confirmed that the FLNC c.2003A>G variant produced an mRNA lacking the terminal 5 bp of exon 12, consistent with the SpliceAI prediction (Fig. 1H). Whereas the 215-bp SEME construct restored the expected in-frame FLNC sequence, the 97-bp SEME construct exhibited partial intron retention extending from FLNC exon 12 to the SEME 5′ splice-site region, resulting in aberrant insertion of an additional 107 nucleotides into the transcript. These findings indicate that insufficient intronic spacing impairs proper splice-site recognition and further identify the 215-bp SEME construct as the most effective configuration for accurate microexon incorporation and restoration of the FLNC reading frame in vitro.

### SEME corrects *FLNC* aberrant splicing and rescues haploinsufficiency in iPSC-CMs

To verify whether SEME can normalize the FLNC c.2003A>G variant in a cardiomyocyte context, we utilized CRISPR–Cas9 technology to generate two distinct iPSC lines: one harboring the heterozygous FLNC c.2003A>G mutation (FLNC ^mut^) and another containing both the mutation and the 215 bp SEME cassette integrated into FLNC intron 12 (FLNC ^SEME^). A schematic overview of the CRISPR–Cas9-mediated knock-in strategy used for SEME insertion into FLNC intron 12 is shown in Fig. 2A. Genomic PCR analysis demonstrated successful insertion of the SEME construct at the targeted FLNC locus (Fig. 2B), and Sanger sequencing further confirmed precise monoallelic integration of the SEME sequence into the mutant FLNC allele (Fig. 2C). The engineered iPSC lines were subsequently differentiated into induced pluripotent stem cell-derived cardiomyocytes (iPSC-CMs) for downstream analyses.

**Figure 2.**
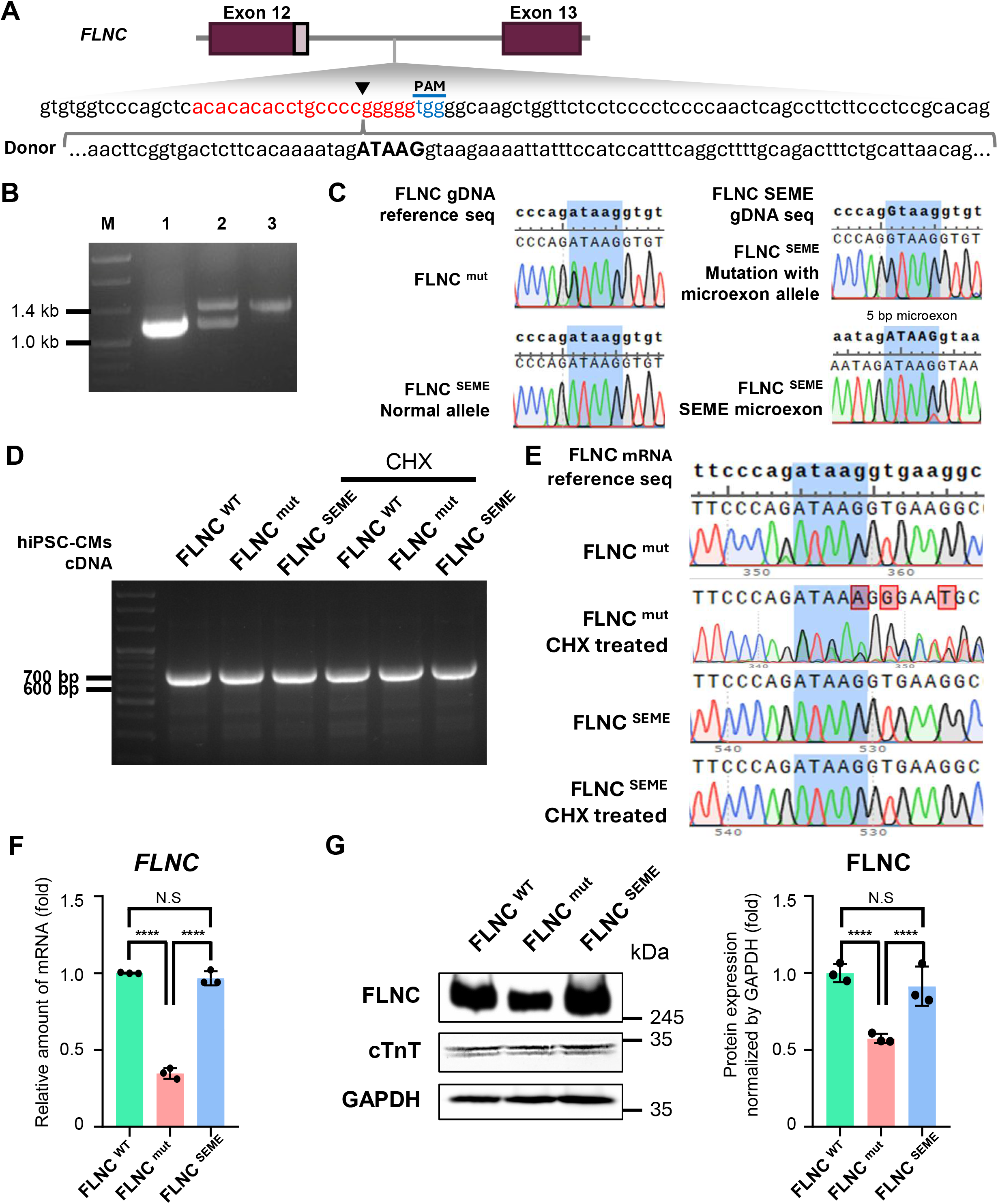
CRISPR-Cas9-mediated SEME integration rescues *FLNC* expression and transcript stability in iPSC-CMs. (A) Schematic diagram of the gRNA targeting sites used to generate the FLNC ^SEME^ cell line. (B) Genotyping of SEME-knock-in iPSCs. Lane 1,2 and 3 represent WT iPSC, FLNC ^SEME^ iPSCs, and the amplified donor template plasmid, respectively.(C) Genomic validation of targeted SEME integration via Sanger sequencing. Representative chromatograms showing the *FLNC* mutations site and the successfully integrated SEME sequence in FLNC ^mut^ and FLNC ^SEME^ iPSC-CMs. The blue-shaded region indicated the 5-nucleotide sequence disrupted by the splicing mutation, which is functionally compensated for by the integration of the synthetic 5 bp microexon. (D) Evaluation of *FLNC* transcript profile via RT-PCR. (E) Transcriptional sequence analysis of splicing rescue. Sanger sequencing chromatograms of FLNC cDNA transcripts before and after CHX treatment in the generated iPSC-CMs. The blue-shaded region highlights the 5-nucleotide deletion site. (F) and (G) Concomitant reduction in mutant lines and rescue of *FLNC* transcript and protein expression. qRT-PCR and immunoblot analyses demonstrated a robust, parallel decrease in both *FLNC* mRNA and filamin C protein level in FLNC ^mut^-CMs. Protein bands were quantified via ImageJ and all expression metrics were normalized to GAPDH. Quantitative data are presented as the mean ±S.D. from three independent experiments. N.S., not significant; **** *p* <0.0001 by one-way ANOVA followed by Tukey’s post-hoc test.

We next examined whether SEME-mediated splicing correction in iPSC-derived cardiomyocytes (iPSC-CMs) recapitulated the splicing patterns observed in the minigene assays. Agarose gel analysis of FLNC cDNA demonstrated no evidence of intron retention in FLNC ^SEME^-CMs, including under 10 µM cycloheximide (CHX) treatment for 4 hours. (Fig. 2D). Under basal conditions, only the wild-type FLNC transcript was detected across all groups. However, following CHX treatment, FLNC ^mut^-CMs exhibited the same aberrantly spliced transcript identified in the minigene assay, whereas no aberrant transcript was detected in FLNC ^SEME^-CMs (Fig. 2E). Consistently, RNA-sequencing coverage showed reduced read depth over the terminal five nucleotides of exon 12 in FLNC ^mut^-CMs, whereas FLNC ^SEME^-CMs exhibited a coverage pattern comparable to wild-type controls (Fig. S2). Sequence alignment of the reconstructed candidate transcript MSTRG.27911.2 supported a net five-nucleotide deletion (ATAAG) at the exon 12 boundary, with CAG microhomology accounting for the apparent exon 13 extension (Fig. S3A). Canonical junction efficiency was 90.86 ± 7.70% in wild-type, 74.24 ± 6.82% in FLNC ^mut^-, and 88.86 ± 0.86% in FLNC ^SEME^-CMs (mean ± SD; n = 3 biological replicates/group). The reduction in mutant cells relative to wild type reached nominal significance (two-sided Welch *p* = 0.0497), whereas the increase following SEME correction did not (*p* = 0.0634; Fig. S3B). Because RNA sequencing was performed without CHX, these measurements reflect steady-state RNA levels and exclude transcripts already degraded by nonsense-mediated decay (NMD), potentially underestimating aberrant transcript production. MSTRG.27911.2 abundance ranged from 7.698 to 217.662 TPM in mutant cells, compared with near-zero levels in wild-type (0–0.081 TPM) and FLNC SEME-CMs (0–0.011 TPM; Fig. S3C). Together, these findings support improved canonical splicing and reduced abundance of the mutant-associated transcript following SEME integration.

Quantitative analysis further revealed that FLNC mRNA expression was markedly reduced in FLNC ^mut^-CMs but restored in FLNC ^SEME^-CMs (Fig. 2F). Similarly, filamin C protein expression was substantially decreased in FLNC ^mut^-CMs and recovered following SEME integration (Fig. 2G). Collectively, these findings demonstrate that SEME-mediated microexon incorporation efficiently corrects FLNC aberrant splicing and rescues haploinsufficiency in iPSC-CMs.

### SEME-mediated correction normalizes sarcomeric organization and calcium handling in FLNC-mutant iPSC-CMs

To evaluate sarcomeric organization in gene-edited iPSC-CMs, immunocytochemistry was performed targeting titin and α-actinin. Both FLNC ^WT^ and FLNC ^SEME^ – CMs exhibited an elongated morphology characteristic of mature cardiomyocytes. Conversely, FLNC ^mut^ – CMs manifested a hypertrophic, rounded phenotype (Fig. S4A). Furthermore, quantitative analysis revealed significantly shortened sarcomere length in FLNC mut-CMs relative to both FLNC WT-CMs and FLNC SEME-CMs, indicating phenotypic rescue following SEME-mediated correction.

To assess calcium handling, we performed a calcium transient assay for in situ monitoring of cytosolic calcium fluctuations. Calcium imaging revealed distinct abnormalities in FLNC ^mut^ – CMs which displayed narrower and more frequent calcium transients compared to FLNC ^WT^ – CMs (Fig. S4B). Quantitative analysis showed that systolic calcium levels and amplitudes in FLNC ^mut^ – CMs exhibited were approximately 13% lower than those in FLNC ^WT^ – CMs (Fig. S4C). Furthermore, FLNC ^mut^ – CMs exhibited significantly faster decay kinetics, with tau and time-to-baseline 50% and 80% values reduced. These results indicate an increasingly divergent reuptake profile as the repolarization phase progressed. Crucially, FLNC ^SEME^ – CMs showed no statistically significant differences from FLNC ^WT^ -CMs across all calcium handling parameters, except for firing frequency. Taken together, these findings demonstrate that SEME-mediated gene correction effectively restore structural integrity, reverses pathological calcium-handling abnormalities, and re-establishes functional homeostasis in FLNC ^mut^ – CMs.

## Discussion

In this study, we developed the SEME, a programmable transcript-reconstruction platform designed to overcome key limitations of conventional exon-targeted therapeutic strategies. Whereas most existing genome-engineering approaches rely on direct modification or removal of protein-coding exons, SEME instead utilizes intronic regions as permissive engineering sites to introduce synthetic microexons through endogenous splicing mechanisms. This strategy enables restoration of disrupted translational reading frames while minimizing direct alteration of native coding sequences.

Exon-skipping approaches can restore disrupted reading frames but necessarily remove coding sequence and, in some cases, require the exclusion of additional exons beyond the mutation-containing exon. In contrast, SEME enables programmable microexon splice-in and can preserve flanking exons that would otherwise need to be skipped. Intronic placement may provide an additional fidelity advantage by preventing local editing byproducts from being transmitted to mature transcripts. Previous work using CRISPR-mediated insertion of exon demonstrated that small insertion–deletion mutations generated at intronic insertion junctions were largely removed during pre-mRNA splicing, resulting in more than 98% correctly processed mRNA products (*21*). This finding provides precedent that intronic editing can buffer mature transcripts from junctional indels that would be disruptive if introduced directly into protein-coding exons. By combining this potential buffering effect with programmable microexon incorporation, SEME may enable transcript reconstruction while minimizing unnecessary exon loss and direct disruption of coding sequences.

The therapeutic potential of SEME was first demonstrated in FLNC-splicing variant iPSC-CMs. FLNC encodes filamin C, a critical cytoskeletal protein in cardiomyocytes, and loss-of-function variants in FLNC are strongly associated with dilated, hypertrophic, and arrhythmogenic cardiomyopathies (*22-24*). FLNC intron 12 spans only 90 bp, providing a substantially more compact splicing environment for microexon incorporation. Despite the compact intronic architecture of FLNC, SEME maintained robust exon-definition activity and enabled accurate microexon incorporation through endogenous splicing mechanisms. Importantly, this molecular correction translated into substantial phenotypic rescue, including restoration of filamin C expression together with normalization of sarcomeric organization and calcium-handling abnormalities. These findings suggest that the SEME architecture is not restricted to its endogenous genomic locus and may be broadly applicable across diverse genes with distinct intronic environments.

Several limitations should be acknowledged. First, SEME was evaluated using engineered isogenic iPSC models rather than patient-derived cells, and only one FLNC variant was examined. Its applicability to other variants, genes, cell types, and intronic environments remain to be established. Second, all experiments were performed in vitro. The efficiency, tissue specificity, durability, immunogenicity, and feasibility of in vivo delivery were not assessed. Third, genomic insertion of SEME requires nuclease-mediated editing and may therefore produce unintended local repair outcomes, off-target alterations, large genomic rearrangements, or clonal selection. Comprehensive genomic safety analyses and validation across independently derived clones will be required. Fourth, the introduction of synthetic splice sites may affect local or transcriptome-wide splicing, and these potential consequences require systematic evaluation. Finally, the amount and sequence of coding information that can be incorporated through SEME are likely to be constrained by microexon size, local splice-regulatory architecture, and tissue-specific splicing activity. The functional consequences of the non-native protein sequences generated by exon replacement must also be evaluated for each application.

Despite these limitations, ongoing advances in genome-engineering technologies are likely to substantially expand the translational potential of SEME. Improvements in genome delivery systems, allele-specific targeting strategies, and precise small-fragment insertion technologies may enable more efficient and selective in vivo implementation of programmable transcript reconstruction. Importantly, SEME may be particularly advantageous in therapeutic settings where preservation of overall protein function is more critical than exact restoration of the native amino acid sequence, provided that functionally tolerated microexons can maintain structural integrity and protein activity.

Collectively, our findings establish programmable microexon incorporation as a proof-of-concept strategy for transcript reconstruction. SEME enabled restoration of sequence lost through aberrant FLNC splicing. These results support further investigation of splicing-guided transcript reconstruction as a complementary approach to direct variant correction and conventional exon skipping.

## Acknowledgement

This research was supported by funding from NIH/NHLBI (R01HL171205) and MSCRF (2024-MSCRFD-6362, 2026-MSCRFL-6569). This research was also supported by a grant of the Korea Health Technology R&D Project through the Korea Health Industry Development Institute (KHIDI), funded by the Ministry of Health & Welfare, Republic of Korea (HI22C0198)

