## supplemental materials for "SEME enables programmable transcript reconstruction through synthetic microexon incorporation"

**Affiliations:**

### Supplementary figure legends

#### Supplementary figure S1. Optimizing SEME flanking sequence length for microexon splicing using AlphaGenome

Predicted splicing profiles for the FLNC c.2003A>G variant and various SEME constructs of differing lengths within *FLNC* intron 12. WT *FLNC* splicing was predicted at positions consistent with the human reference sequence (top panel, *FLNC* exon 11 – intron 24). In the patient variant, the canonical splice donor site of exon 12 was abolished, and a cryptic splice donor was predicted to occur 5 bp upstream at the native site (second panel). A SEME construct incorporating the minimum essential sequences, 67 bp of 3'ss and 6 bp of 5'ss, yielded a microexon splice-in score of approximately 0.5 (third panel). However, extending the 5'ss to 25 bp resulted in splice-in scores approaching the maximum threshold. While the microexon splice-in scores remained at the maximum across all extended SEME constructs, intron sequences exceeding 1 kbp in length led to the emergence of cryptic splice acceptor and donor sites within the intronic region. Red arrows indicate the position of the FLNC c.2003A>G variant; blue arrows denote the position of the SEME-derived microexon.

#### Supplementary figure S2. Sashimi plot analysis of *FLNC* exon 12-13 splicing pattern in *FLNC* $^{WT}$, *FLNC* $^{mut}$, and *FLNC* $^{SEME}$ -CMs.

Splicing profiles of the *FLNC* exon 12-13 region identified by RNA-seq in *FLNC* $^{WT}$, *FLNC* $^{mut}$, and *FLNC* $^{SEME}$ – CMs. In *FLNC* $^{mut}$ – CMs, a marked reduction in read depth was observed at the last 5 bp of exon 12, with aberrant splicing. In contrast, the splicing architecture of *FLNC* $^{SEME}$ – CMs was indistinguishable from that of *FLNC* $^{WT}$ – CMs, demonstrating the successful restoration of the canonical splicing pattern by SEME integration.

#### Supplementary figure S3. A mutant-associated FLNC transcript consistent with a five-nucleotide deletion is not detected after SEME correction.

(A) Alignment of FLNC-201 and the assembled transcript MSTRG.27911.2 at the exon 12–13 junction. Although the merged GTF assigns a repeated CAG motif to the 5′ end of exon 13, alignment of the spliced sequences favors deletion of ATAAG from the 3′ end of exon 12. The apparent three-nucleotide extension of exon 13 is therefore consistent with boundary ambiguity arising from CAG microhomology, rather than an independent insertion. (B) Canonical exon 12–13 junction efficiency in wild-type, FLNC-mutant, and SEME-corrected cardiomyocytes. (C) MSTRG.27911.2 abundance in the same groups. Points represent biological replicates (n = 3/group); horizontal lines indicate group means, and error bars in B indicate SD.

**Supplementary figure S4. Intracellular structure and calcium handling characterization of WT, mutant, and SEME-corrected iPSC-CMs.** (A) Sarcomeric architecture of FLNC $^{WT}$, FLNC $^{mut}$, and FLNC $^{SEME}$ – CMs. FLNC $^{WT}$ and FLNC $^{SEME}$ -CMs exhibited an elongated morphology typical of mature cardiomyocytes, characterized by a highly organized, linear alignment of sarcomeres. In contrast, FLNC $^{mut}$-CMs displayed an enlarged, rounded cellular phenotype, featuring regions with disorganized, orthogonally arranged sarcomeric structures. Furthermore, the sarcomere length was significantly shortened in FLNC $^{mut}$ – CMs. (B) Representative calcium transient traces. Fluorescence intensity profiles capturing intracellular calcium dynamics in iPSC-CMs. FLNC $^{mut}$-CMs manifested a narrower calcium pulse width and an accelerated firing frequency compared to WT controls, whereas FLNC $^{SEME}$-CMs displayed completely restored pulse widths and kinetic frequencies. (C) Statistical analysis of calcium transient kinetics. Comprehensive evaluation of intracellular calcium handling parameters. FLNC $^{mut}$-CMs exhibited a significant decrease in systolic calcium amplitude but an elevated firing frequency. Furthermore, the mutant cells displayed significantly accelerated clearance kinetics across all reuptake parameters. Each data point represents a single-cell measurement. (n = 25). Data was presented as the mean ±S.D. from three independent experiments. N.S., not significant; *$p$<0.05, **$p$<0.01, ***$p$<0.001, ****$p$<0.0001 by one-way ANOVA followed by Tukey's post-hoc test.

**Table S1. List of oligonucleotide sequences used for PCR and Sanger Sequencing.**

| Name | Sequence (5' to 3') |
|---|---|
| **Primers for minigene construction** | |
| FLNC minigene | F : TTGCTATAGCACGAGCTCATTTTGGGGTCCTGGTTTGGAGAC<br>R : GTGAGGGTCTCCTGCTATTTCGTACACCTTTACCCGCTCG |
| FLNC mX5 67 bp | F: CTCACACACACCTGCCCCTTTGATTTTTGTTTCTTTTTTTTCCATGTCTGTCCTGTCTGAACTTCGGTGACTCTTCAC<br>R: CCAGCTTGCCCCACCCCCAAATGGATGGAAATAATTTCTTACCTTATCTATTTTGTGAAGAGTCACCGAAGTTC |
| FLNC patient minigene mutagenesis | F : CCCACCTGACTGCTTCCCAGGTAAGGTGTGGTCCCAGCTCA<br>R : TGAGCTGGGACCACACCTTACCTGGGAAGCAGTCAGGTGGG |
| FLNC mX5 215 bp | F : CTCACACACACCTGCCCCGTGTGAGGTTGCAGTAACCTAATAAGAC<br>R : CCAGCTTGCCCCACCCCCCGGCTAGAAAGGCTCTCATATGCA |
| FLNC mX5 447 bp | F : CTCACACACACCTGCCCCTTCTATTGGACAGTTCTGCTCTAAGG<br>R : CCAGCTTGCCCCACCCCCTGTCTAGGCTCCCCAGGACA |
| FLNC mX5 mutagenesis | F : CTTCGGTGACTCTTCACAAAATAGATAAGGTAAGAAAATTATTTCCATCCAT<br>R : ATGGATGGAAATAATTTTCTTACCTTATCTATTTTGTGAAGAGTCACCGAAG |
| **Sequencing primer sequences** | |
| Genomic FLNC | F : CATGCTGTCCTGTCTAGGCCATC<br>R : GGACACCAGTCTGAGAAACTGACTTG |
| cDNA FLNC | F : TGCCTGGGAAGTATGTG<br>R : AAGTAGGTGGGCTCATT |
| 5'UTR from ALDOA | F : CTCCGTCTGGATTTCCAAGGAA |
| EGFP | R : GAACTTGTGGCCGTTTACGT |
| FLNC mut dsDonor | F : TGGGCAAGTCAGCCGATTTTGTGG<br>R : TCACCCAAGCAAAGTTCATAGCTGTGG |
| FLNC SEME dsDonor | F : TATGGCCAGTGGTTCTGACTC<br>R : TGCCCTCCCTCAAGGCTCAT |

**Table S2. List of oligonucleotide sequences used for qRT-PCR.**

| Name | Sequence (5’ to 3’) |
|---|---|
| FLNC | F : CCTATGGGCCTGGCATCGAG<br>R : GGGAACCACCTTAGCCTCCT |
| GAPDH | F : TGCACCACCAACTGCTTAGC<br>R : GGCATGGACTGTGGTCATGAG |

**Table S3. List of used antibodies.**

| Name | Supplier | Cat No. | Dilution ratio |
|---|---|---|---|
| **Western blot** | | | |
| Filamin C | Novus Biologicals | NBP3-04857 | 1:1000 |
| GAPDH | Cell Signaling Technology | 2118 | 1:1000 |
| Cardiac Troponin T | Invitrogen | MA5-12960 | 1:1000 |
| Goat anti-Mouse IgG (H+L) Cross-Adsorbed Secondary Antibody, HRP | Invitrogen | A16072 | 1:5000 |
| Goat anti-Rabbit IgG (H+L) Cross-Adsorbed Secondary Antibody, HRP | Invitrogen | G21234 | 1:5000 |
| **Immunocytochemistry** | | | |
| Titin | Proteintech | 27867-1-AP | 1:100 |
| α-actinin | Thermo Fisher Scientific | A7811 | 1:100 |
| Chicken anti-Mouse IgG (H+L) Cross-Adsorbed Secondary Antibody, Alexa Fluor™ 488 | Invitrogen | A21200 | 1:500 |
| Goat anti-Rabbit IgG (H+L) Highly Cross-Adsorbed Secondary Antibody, Alexa Fluor™ 546 | Invitrogen | A11035 | 1:500 |

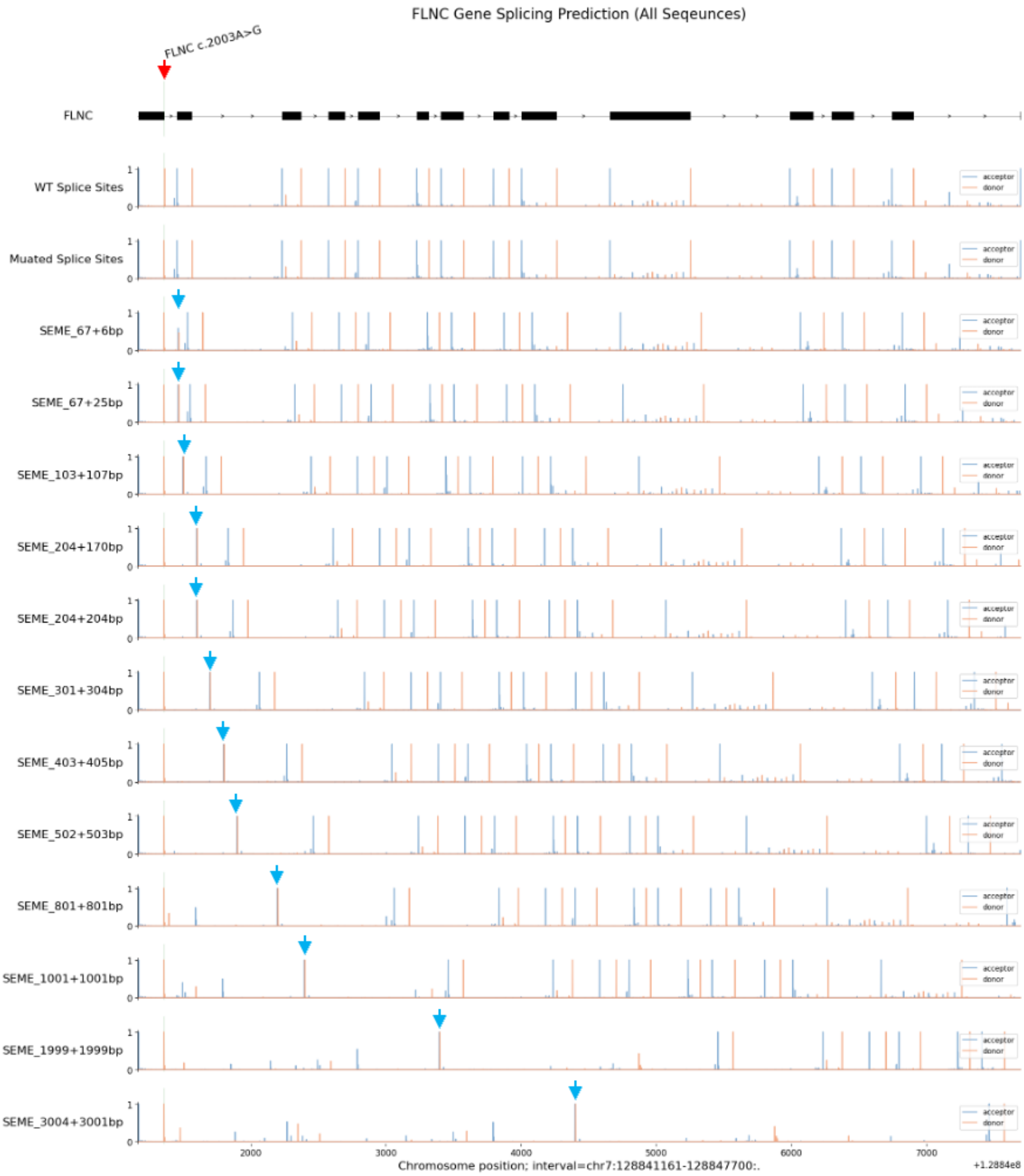


**Supplementary figure S1. Optimizing SEME flanking sequence length for microexon splicing using AlphaGenome**

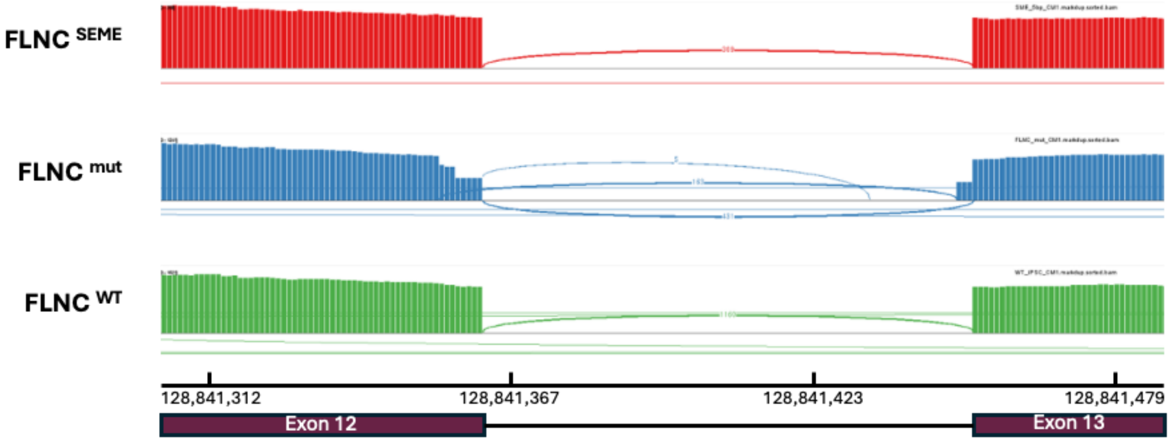


**Supplementary figure S2. Sashimi plot analysis of *FLNC* exon 12-13 splicing pattern in *FLNC* $^{WT}$, *FLNC* $^{mut}$, and *FLNC* $^{SEME}$ -CMs**

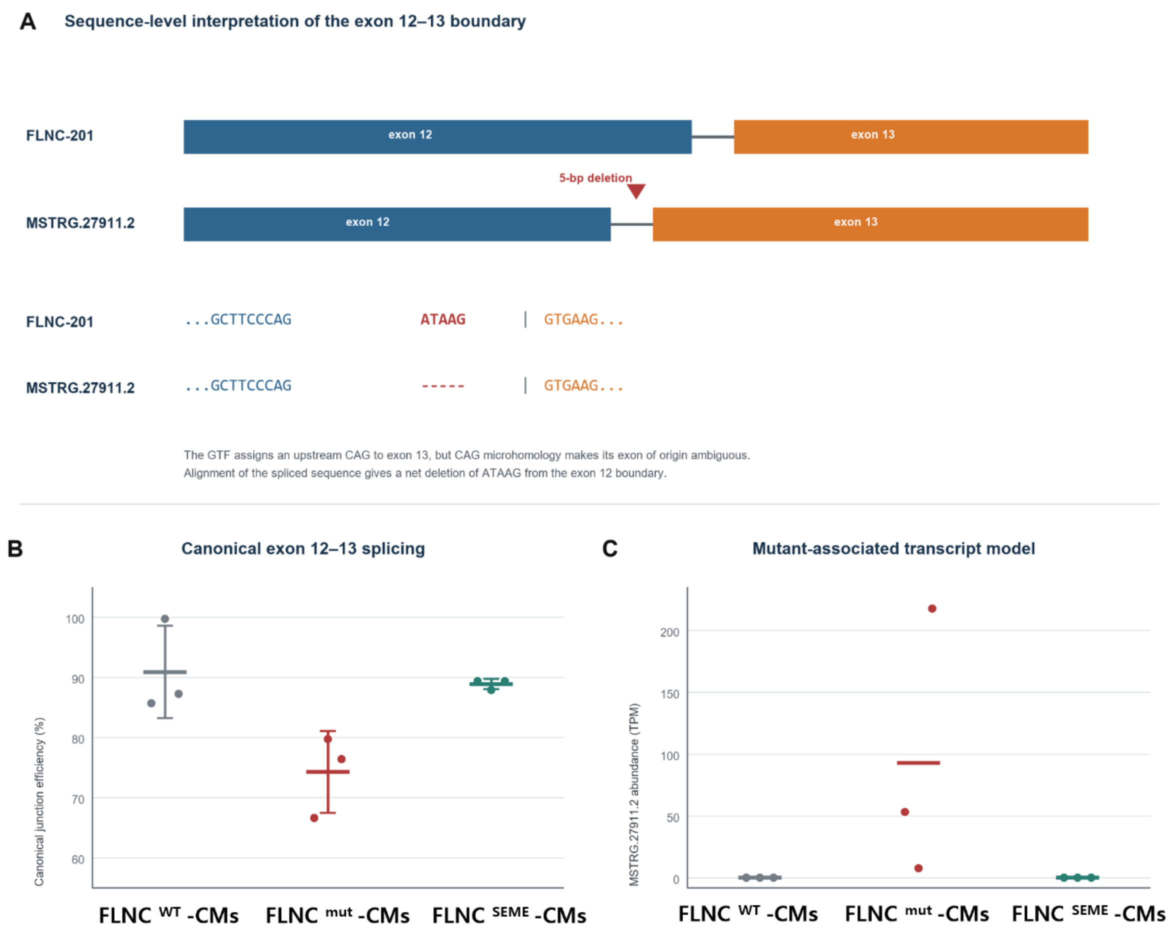


**Supplementary figure S3. A mutant-associated FLNC transcript consistent with a five-nucleotide deletion is not detected after SEME correction.**

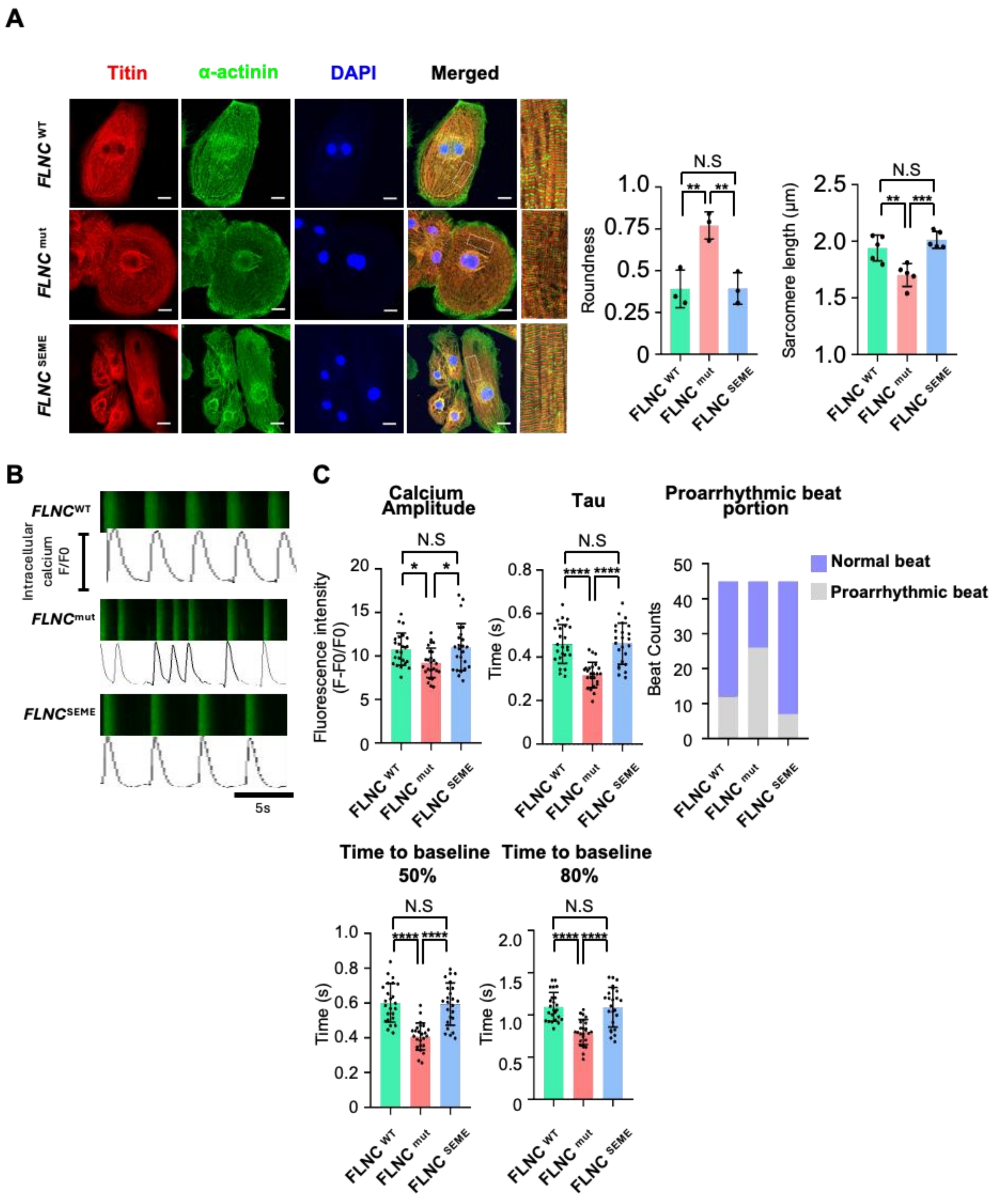


**Supplementary figure S4. Intracellular structure and calcium handling characterization of WT, mutant, and SEME-corrected iPSC-CMs.**
